# StrainSeeker: fast identification of bacterial strains from unassembled sequencing reads using user-provided guide trees

**DOI:** 10.1101/040261

**Authors:** Märt Roosaare, Mihkel Vaher, Lauris Kaplinski, Märt Möls, Reidar Andreson, Maarja Lepamets, Triinu Kõressaar, Paul Naaber, Siiri Kõljalg, Maido Remm

**Affiliations:** Department of Bioinformatics, IMCB, University of Tartu, Riia 23, EE-51010 Tartu, Estonia; Estonian Biocentre, Riia 23, EE-51010 Tartu, Estonia; Institute of Mathematical Statistics, University of Tartu, Liivi 2, EE-50409 Tartu, Estonia; Department of Microbiology, University of Tartu, Ravila 19, EE-50411 Tartu, Estonia; synlab Eesti, Väike-Paala 1, EE-11415 Tallinn, Estonia; United Laboratories, Tartu University Clinics, L. Puusepa 1 a, EE-50406 Tartu, Estonia

**Keywords:** k-mer, microbiome, strain identification, species identification, diagnostics

## Abstract

**Background:** Fast, accurate and high-throughput detection of bacteria is in great demand. The present work was conducted to investigate the possibility of identifying both known and unknown bacterial strains from unassembled next-generation sequencing reads using custom-made guide trees.

**Results:** A program named StrainSeeker was developed that constructs a list of specific k-mers for each node of any given Newick-format tree and enables rapid identification of bacterial genomes within minutes. StrainSeeker has been tested and shown to successfully identify *Escherichia coli* strains from mixed samples in less than 5 minutes. StrainSeeker can also identify bacterial strains from highly diverse metagenomics samples. StrainSeeker is available at http://bioinfo.ut.ee/strainseeker.

**Conclusions:** Our novel approach can be useful for both clinical diagnostics and research laboratories because novel bacterial strains are constantly emerging and their fast and accurate detection is very important.

## Background

Pathogenic bacteria represent a considerable danger for human health worldwide. Often, it is hard to pinpoint the exact strain or species causing the disease in a timely manner using conventional methods such as culturing, which can take at least a full day plus analysis [1]. Moreover, it is very important to differentiate between bacterial strains because they can have vastly different effects on their host. A well-known example is *Escherichia coli* sp., which contains some strains such as *E. coli* O157:H7 [2] and *E. coli* EC958 [3] that are considerably more virulent than others. Methods and databases based on 16S RNA sequences such as the well-known Ribosomal Database Project [4] have been used to characterize diverse bacterial populations, but they do not have the necessary resolution to identify bacteria at the strain level [1, 2]. In recent years, matrix-assisted laser desorption/ionization time-of-flight mass spectrometry has been used to quickly and cheaply identify bacterial colonies [5], but for strain-level identification it requires very precise, manually crafted databases for each species which, to a large extent, are not available today. One solution is to use whole genome sequencing (WGS) directly on clinical samples to obtain the data as quickly as possible, thereby avoiding the time spent on cultivating bacterial isolates. WGS can also detect pathogens that grow very slowly or demand very specific conditions and therefore are difficult to cultivate [1]. However, WGS generates extremely large amounts of data (ranging typically from 500 million (M) base pairs (bp) to 3000 Mbp [6] per sequenced sample), which makes information processing challenging. State-of-the-art bioinformatic tools such as Kraken [7], CLARK [8], CoMeta [9] and KmerFinder [1] are based on the detection of short DNA oligomers with length *k* (k-mers). Kraken and CoMeta identify each of the sequence reads separately using the National Center for Biotechnology Information (NCBI) taxonomy tree, counting the hits to each of the taxons on the tree and finding the branch with the most total hits. CLARK also identifies each of the reads, but instead of using a tree, it is based on a flat, user-defined database. KmerFinder counts the k-mer hits to each of its database strains and computes the significance of the matches.

A strain-level tree for all bacteria would also help to avoid controversies such as the case of *E. coli* and *Shigella* sp., where *Shigella* strains are phylogenetically similar to *E. coli* but are different species according to NCBI taxonomy [10].

We present StrainSeeker, a program for the strain-level identification of bacteria from raw sequencing reads either from an isolate or a metagenomic sample. Unlike other k-mer based classifiers, StrainSeeker does not classify each read separately but analyzes all of the k-mers in the sample together. The program uses a guide tree to represent relationships between different bacteria down to the strain level, not being tied to existing taxonomic systems such as the NCBI taxonomy.

## Results and discussion

### Overview of the method

StrainSeeker consists of two main steps: database building (or downloading a pre-built database) and detection of strains from the sequenced sample.

#### 1. Creating the k-mer database

Before StrainSeeker can be used to identify bacteria, the database of specific k-mers needs to be built or downloaded. Lists of the k-mer database can be pre-built for all known bacterial genomes or created by the end-users for a custom set of strains. For this, the user has to provide genomic DNA sequences (in the form of full genomes or assembled contigs) from a set of different strains and a guide tree that includes all of the provided strains. The tree can be built using any method preferred by the user as long as it is a binary tree in Newick format. Then, specific k-mer lists for each node and strain on the tree are calculated using the tree and genome sequences, thereby creating a hierarchical structure of k-mer lists (the “database”).

#### 2. Identification of bacterial strains from the sample

StrainSeeker converts raw sequence reads (FASTA or FASTQ) of the sample to k-mers and compares them with the k-mer database to detect known and/or new strains.

### Advantages and limitations of guide tree-based strain detection

Bacteria evolve fast. Therefore, it is necessary to detect emerging strains because they can have very different phenotypes compared to their relatives. To solve this problem, we use a guide tree that allows us to also detect unknown strains that are close relatives to strains included in the database. We decided not to use NCBI taxonomy because it did not contain strain-level relationships, making it unsuitable for the detection of closely related strains. Also, in the NCBI tree some taxons are not monophyletic e.g. the *Shigella/Escherichia* branch. In the case of highly similar strains, all of the k-mers that are specific to both of them are moved up to their last common ancestor, leaving very few strain-specific k-mers. With the help of the guide tree, StrainSeeker is able to detect the last common ancestor on the sub-species level. For the present work, we used an alignment-free k-mer-based distance method similar to the k-mer natural vector method [11, 12] to construct the tree of 2,758 bacterial strains. To test the accuracy of our tree, the *E. coli* sp. and *Shigella* sp. subtree (Figure 1) was compared to two gene-based trees [13, 14] and a k-mer based tree [15], indicating that our alignment-free k-mer-based distance method can be used to accurately represent strain-level relationships. Higher-level relationships on the tree (genus level and above) are irrelevant for StrainSeeker as our database is made of subtrees that mainly consist of a single species. In general, the more inaccurate the tree, the less node-specific k-mers the database will contain, resulting in a flat strain-level database similar to CLARK [8] in an extreme case.

**Figure 1.**
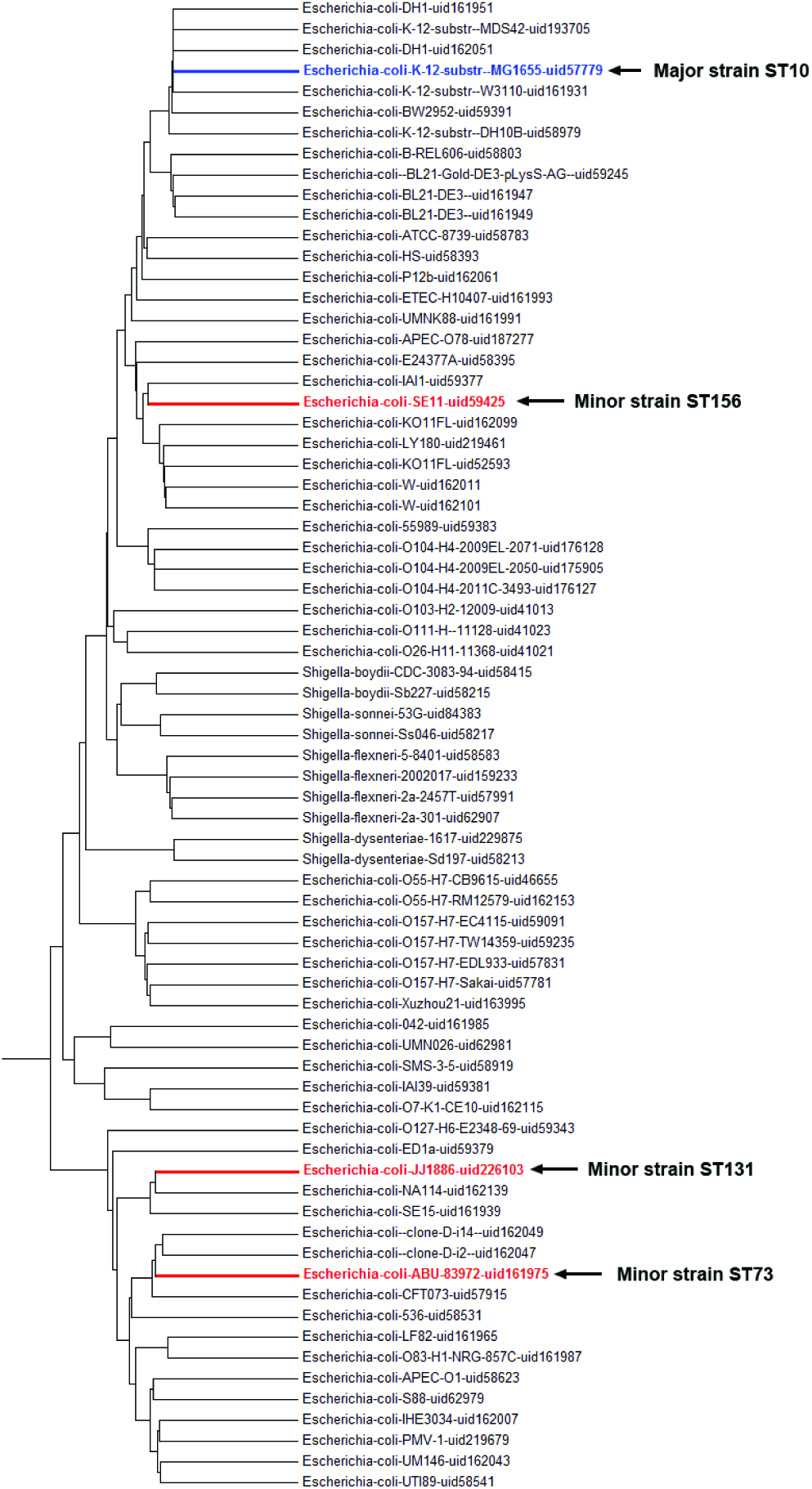
*E. coli sp.* and *Shigella* sp. subtree from our guide tree. The tree is a subtree from the guide tree of 2,758 strains, based on the fraction of shared k-mers. Organism names contain the NCBI RefSeq name and the bioproject identifier. Branches with *E. coli* strains most closely related to the minor strains used in the sensitivity and strain proportion measuring experiments are highlighted in red, the major strain in blue. The tree was constructed in MEGA 6 [16] using the UPGMA method.

### Building the k-mer database

To create the specific k-mer database, the user needs to provide a set of bacterial genomic DNA sequences and a guide tree. The k-mer database is built according to the guide tree structure (Figure 2). The building process starts from the leaves and moves towards the root. To reduce the noise in samples that is caused by the DNA of other, non-bacterial organisms such as human DNA in clinical samples, the user can also provide a list of potential contaminating sequences (the “blacklist”) that is used to eliminate all strain k-mers that are also present in the blacklist. The blacklist itself is not part of the database. The final database contains specific k-mers for each internal and external node (strain) represented in the guide tree and an index file containing the database structure and k-mer counts. In our study, we created the databases with *k=32* using the GenomeTester4 software [17]. Longer k-mers would give only strain-specific k-mers, shorter k-mers would give only node specific k-mers. In our experience, *k=32* is optimal for the strain-level identification of bacteria as it gives an optimal distribution between node-specific k-mers (Figure 2, node1 and node2) and strain-specific k-mers (Figure 2, strain 1-3). Also, we restricted the maximum amount of k-mers in a node or strain to 100,000 as larger amounts did not improve the accuracy (data not shown). The main database was constructed from 2,758 bacterial genomes obtained from the NCBI RefSeq database, its size was approximately 7.8 GB. The blacklist that we used consisted of k-mers from the human genome assembly GRCh38 and all the plasmids of the 2,758 bacteria. The database consisted of many subtrees as very different bacteria apparently have very few or no common k-mers if *k=32*.

**Figure 2.**
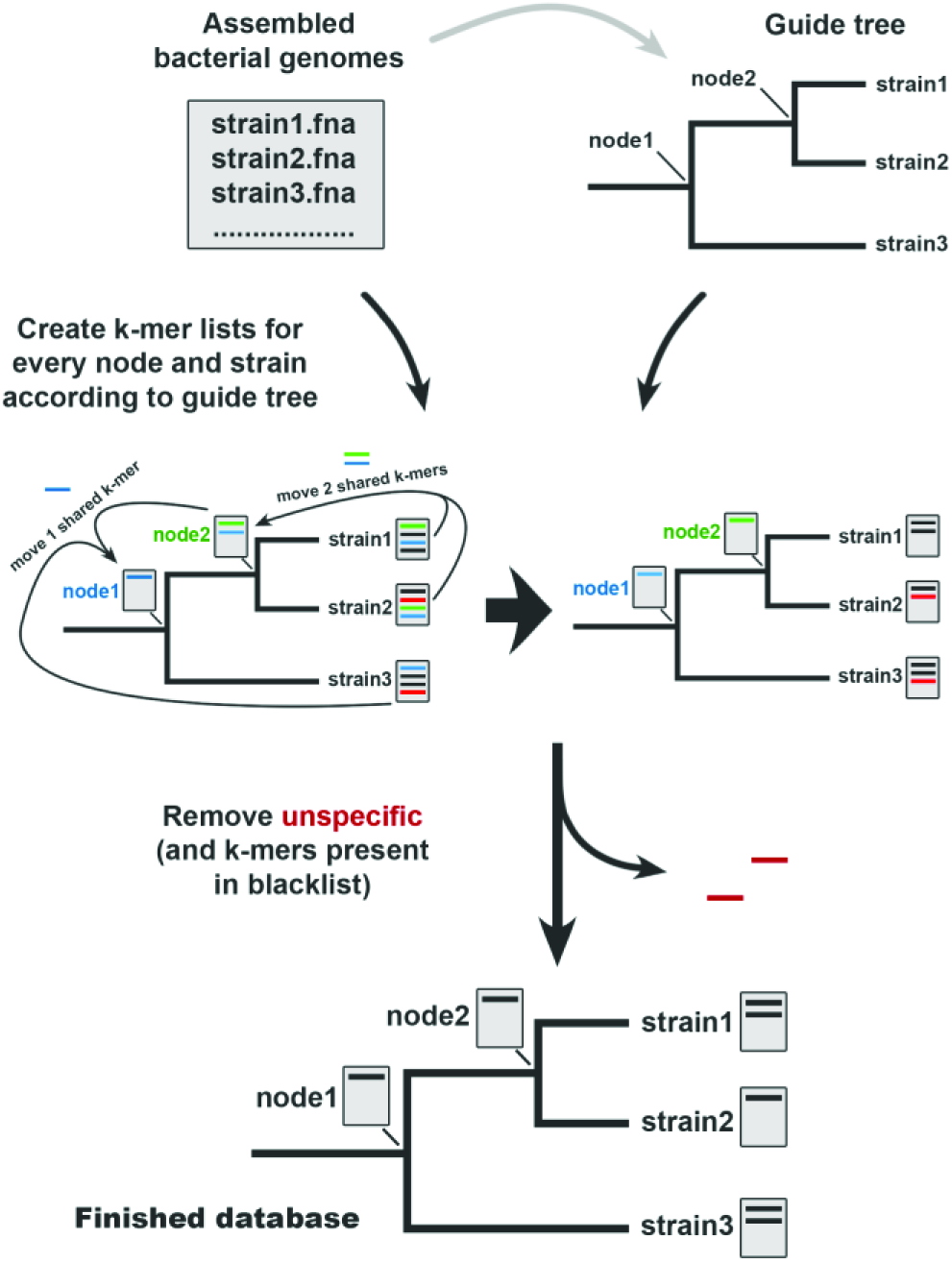
StrainSeeker database building process. Database construction starts with the selection of bacterial strains of interest. The assembled genome of each strain is converted into a k-mer list. Next, a guide tree is created for the database. We used the strain k-mer lists and a k-mer-based method to construct the guide tree but any tree in Newick format can be used. The building process starts from the strain level and moves shared k-mers towards the root. The final step is to eliminate non-specific k-mers that occur in the “blacklist” or in any other nodes. The finished database contains k-mer lists specific to each node and strain and can be used to quickly identify any strain included on the guide tree and the strains related to them.

### Strain identification algorithm

StrainSeeker does not attempt to identify each read separately, but analyzes all the specific k-mers of a strain that are found in the sample. All other k-mer based identifiers tested in a recent publication will try to identify each read [18]. The main advantage of our approach is that we use a ratio (k-mers found in the sample divided by all the queried k-mers; same k-mer is counted only once). Therefore, the decision of calling a strain is based on all the k-mers of the strain whereas other programs have to make this decision for each read, based on a limited number of k-mers. This makes StrainSeeker less vulnerable to errors if there are lots of non-specific k-mers in the sample (due to technical or biological reasons) which, according to the database, are specific.

The core of the search process is the guide tree structure. The search is recursive, starting at the root node of the tree (or subtree) and moving down towards the strains (Figure 3). The percentage of observed k-mers is calculated at each step. This helps to constrain the search space because we can skip all branches where the percentage is too low. We assume that node-specific k-mers are located randomly across a bacterial genome and sequencing covers the genome randomly as well. Therefore, we can infer that we should observe the same proportion of k-mers (O) in each of the nodes on the path from the root node to the strain. Also, the proportion of k-mers in a node is approximately equal to the cumulative proportion of k-mers in the child nodes (E; the exact formula can be found in the methods section). O/E > 1 means that one or both of the child nodes contribute less than the expected amount of k-mers, indicating the presence of a strain that is most similar to the current node but not to any of the sub-nodes or leaves under it (Figure 3, N4). In this case, StrainSeeker will output all of the strains under the current node. *O/E* = 1 indicates that the strain belongs to one of the sub-nodes and the search process will continue (Figure 3, N1, N2 and N3). O/E < 1 indicates sequencing errors or contamination (Figure 3, N5).

**Figure 3.**
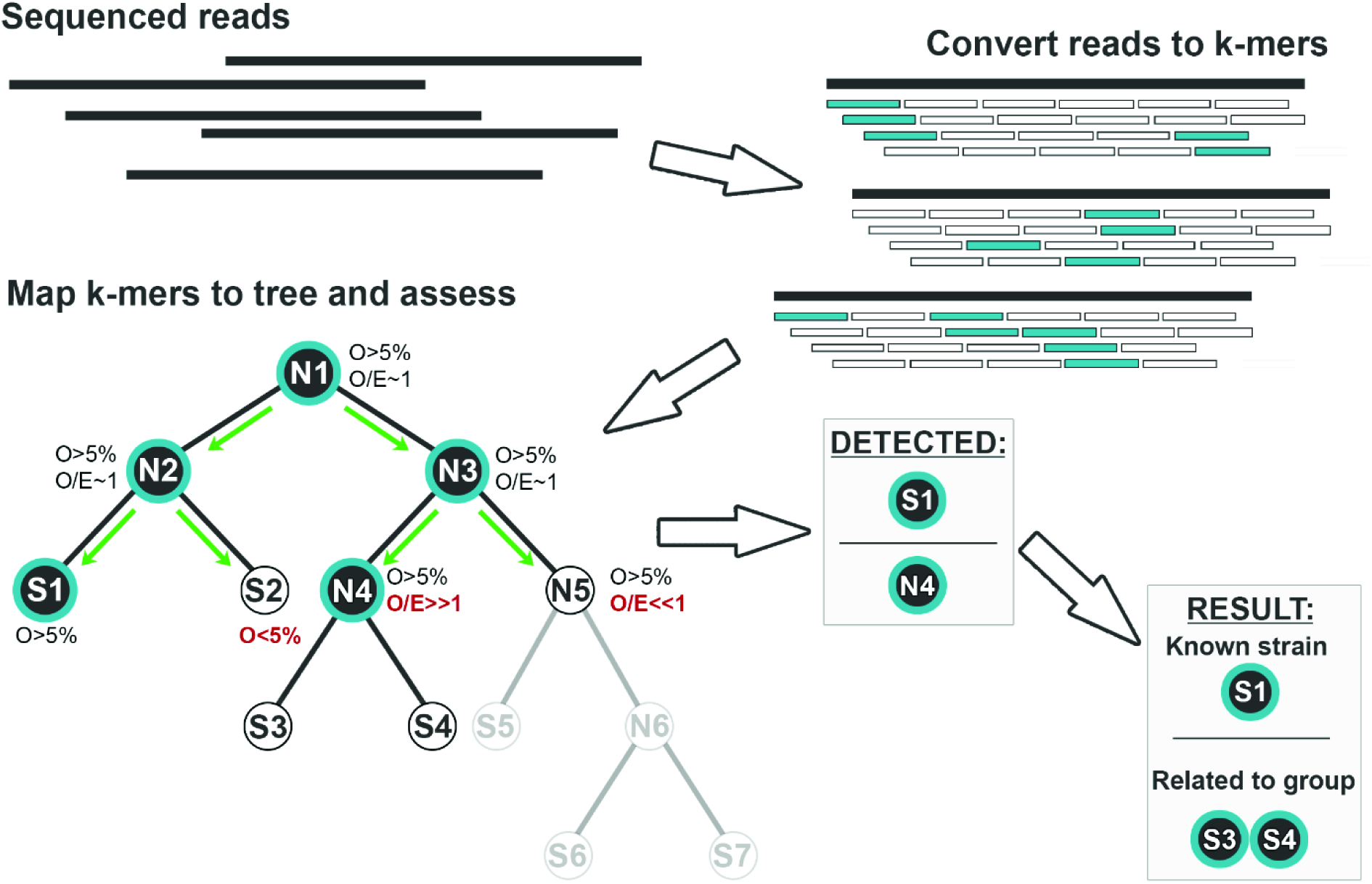
Strain identification process. After the sample is sequenced, the reads are converted into k-mers. The search starts from root (N1) and recursively moves down to the subnodes according to the following criteria. First, the fraction of observed k-mers (O) is calculated. For the identification process to continue along the current branch, *O* has to exceed the cutoff level (5%). Then, an observed/expected (O/E) value is calculated for nodes or the strain is shown as the result for the leaves. The process continues to the subnodes if *O/E = 1*, showing that the current node is present in the sample and there is no branching to unknown nodes and strains. Significantly higher O/E values (N4) indicate the presence of an unknown strain that is most closely related to N4. Single strains are provided (S1) in the output in the case of known strains. Conversely, all strains that are under an internal node (S3, S4) are provided if the strain(s) most closely related to this internal node (N4) is present in the sample. The result above indicates that two strains were found in total: known strain S1 and a strain that is closely related to the known strains S3 and S4.

StrainSeeker will show all strains located under the node closest to the unknown strain as the result (Figure 4). Additionally, relative amounts of all identified strains (relative fractions of each genome in the sample) are shown, calculated using the ratio of specific k-mers found to the total amount of specific k-mers of a strain. This measure represents the actual abundances better than identified reads because organisms with larger genomes generate more reads [19].

**Figure 4.**
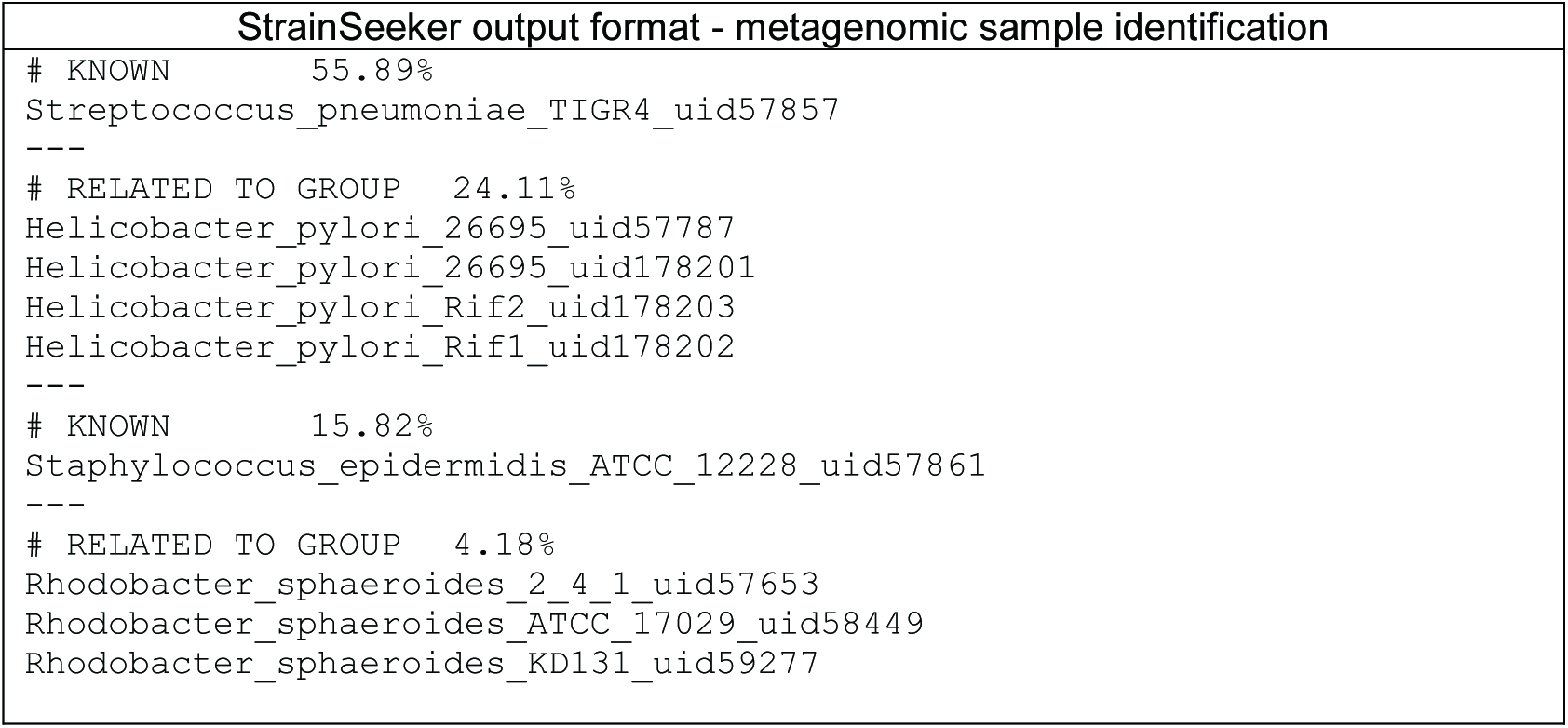
StrainSeeker output format. This is an example of the StrainSeeker output for a metagenomic sample that consisted of four different strains. StrainSeeker will output all separate strains identified in the sample. For known strains, “KNOWN” and the strain name is given (Figure 3, S1). In the case of unknown strains that are similar to a group of strains on the guide tree, “RELATED TO GROUP” along with a list of related strains is given (Figure 3, N4). StrainSeeker will also calculate relative genome frequencies for all of the detected strains.

### Testing the sensitivity of StrainSeeker using samples consisting of similar strains

Biological samples can often contain multiple strains from the same species, which can make it difficult to detect the strain with lower abundance (the “minor” strain). In order to test StrainSeeker’s sensitivity in such a case, we created six artificial test samples for each of the three minor strains, 18 samples in total (Table 1). Every sample consisted of 1 million Illumina 100 bp reads. In each sample, variable amounts of the minor strain were mixed with an *E. coli* ST10 strain (*E. coli* K12 substrain MG1655), which served as the major strain. We used *E. coli* strains with MLST type ST73, ST131 and ST156 isolated from clinical samples as the minor strains. The database used in all the experiments was the one mentioned above, containing 2,758 strains.

**Table 1.**
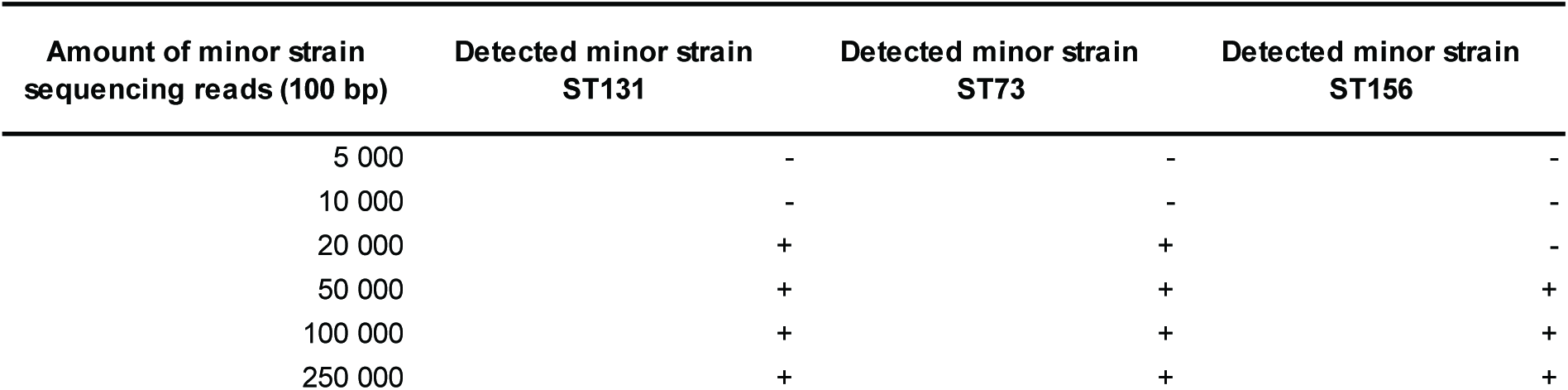
StrainSeeker predictions for the three minor strain proportions in 18 samples, each consisting of 1 million Illumina reads. “+” stands for called, “-” for not called.

| Amount of minor strain<br>sequencing reads (100 bp) | Detected minor strain<br>ST131 | Detected minor strain<br>ST73 | Detected minor strain<br>ST156 |
| --- | --- | --- | --- |
| 5 000 | - | - | - |
| 10 000 | - | - | - |
| 20 000 | + | + | - |
| 50 000 | + | + | + |
| 100 000 | + | + | + |
| 250 000 | + | + | + |

StrainSeeker successfully detected the minor strain when at least 20,000 reads (50,000 in the case of ST156) were present despite the noise caused by the high levels of the major strain. The only ambiguous result given by StrainSeeker was the sample with 10,000 ST73 reads in which ST73 was identified as “Related to” a group of strains that also contained some ST131 strains. This could be due to the low amount of ST73 reads as the results improved with higher read amounts. As StrainSeeker does not classify each of the reads separately, we cannot compare the precision and sensitivity of read identification as presented in other recent papers [7–9].

### Testing StrainSeeker’s accuracy in measuring strain proportions

We wanted to test how accurately StrainSeeker can estimate the ratios of different strains in mixed samples. We used the same 18 samples that were also used in the sensitivity testing. StrainSeeker accurately measured the proportions of *E. coli* ST73 strain in the samples (Figure 5) and predicted lower proportions for the ST131 and ST156 strains. This result could be because the ST73 strain had a more similar strain present in the database than the ST131 and ST156 strains.

**Figure 5.**
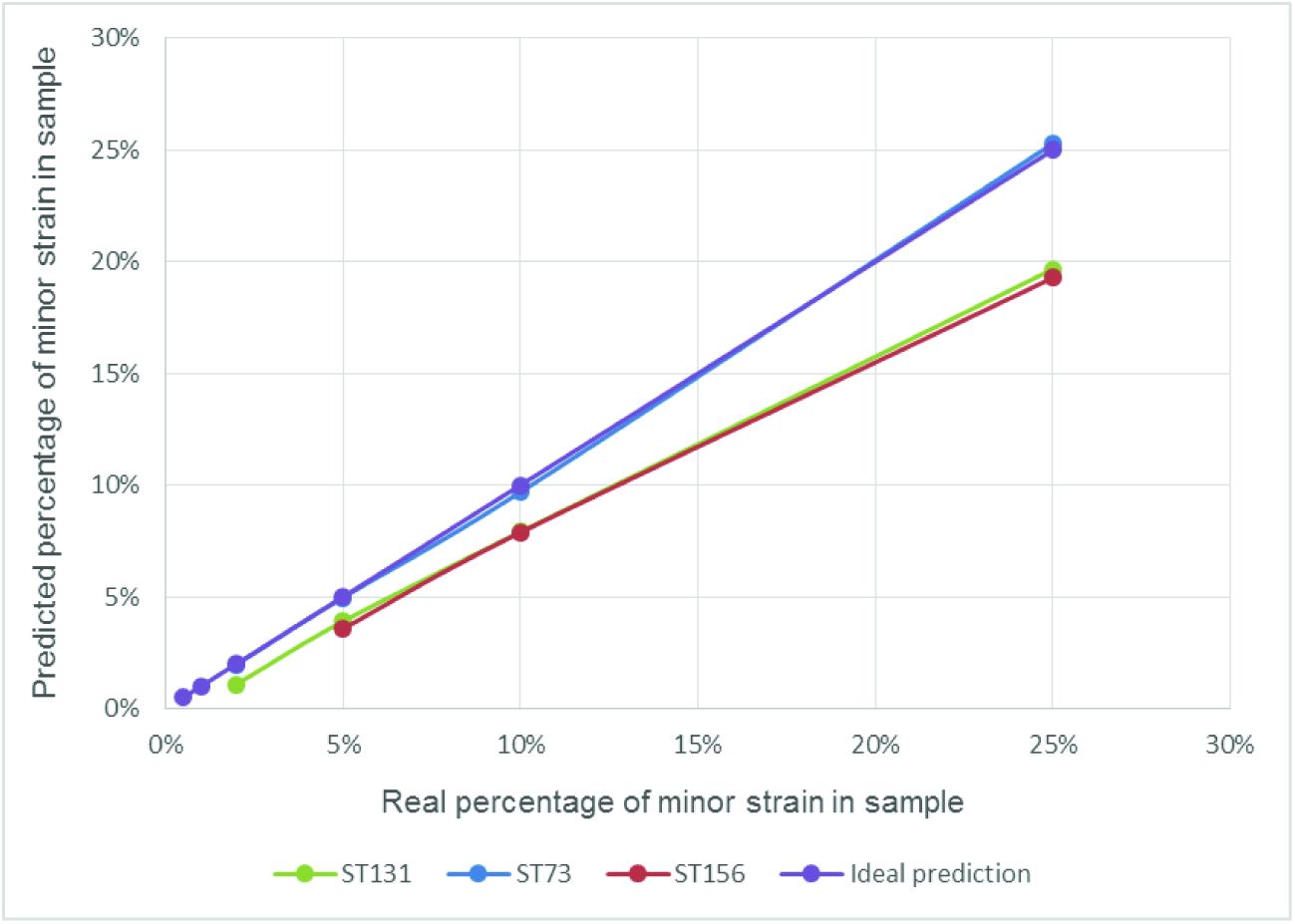
StrainSeeker’s accuracy in assessing the proportions of three minor *E. coli* strains in mixed samples. We created 18 artificial samples (Table 1) of 1 million 100 bp Illumina reads. *E. coli* K12 substrain MG1655 constituted the majority of each sample (75% to 99.5%), minor strains used were *E. coli* ST131, ST73 and ST156 (0.1% to 25%). StrainSeeker was able to detect ST131 and ST73 with a minimum of 20,000 reads and ST156 with a minimum of 50,000 reads. Assessing the proportions of strains in the samples was very accurate for ST73. In case of ST131 and ST156 strains, StrainSeeker systematically predicted lower amounts. This might be because the ST73 strain had a more similar strain present in the database than the ST131 and ST156 strains.

### Testing StrainSeeker’s sensitivity and precision on a metagenomic sample

We wanted to test how well StrainSeeker will perform if the sample contained a diverse population of bacteria instead of just an isolate or a few similar strains. For that, we used a mock community metagenomic sample SRR172902 obtained from the Sequence Read Archive (SRA), which contained 21 bacteria. To make the sample more similar to a real microbiome, which would often contain human DNA as well as bacterial, we added human raw sequencing reads (SRA database, ERR1055492). Final ratio of bacterial reads to human reads in the 4.92 Gbp sample was 1:10, human DNA constituting the majority of the sample.

The sample was identified with StrainSeeker using the 2,758 strain database (results provided in additional data file 2). StrainSeeker correctly identified 20 of the 21 strains, despite the high amount of noise caused by the human DNA. The unidentified bacterium, *Actinomyces odontolyticus*, was not present in our database and therefore could not be detected. StrainSeeker’s sensitivity was 100% and precision 100%. The presence of human DNA does not hinder StrainSeeker, therefore we assume that it can be used to identify bacterial strains from complex samples containing human DNA background.

### Comparing StrainSeeker to other metagenomics analysis tools

To compare StrainSeeker to other popular tools, we selected a dataset that was recently used by a study that thoroughly compared 14 tools [18]. As mentioned before, we cannot compare the precision and sensitivity of read identification, therefore we decided to analyse the amount of genera found in the sample. Genus level was chosen as the other level examined in the study (phylum) was too broad. In our case, false positive means that a genus not present in the sample was found and false negative indicated a genus present in the sample that was not called by StrainSeeker. We used one replicate from each of the two samples (set A1 and set B1), calculated metrics are the averages of both sets. StrainSeeker’s run time was 12.3 minutes, with genus-level sensitivity 0.968 and precision 0.995. Most of the false negatives were due to these strains missing from our database. StrainSeeker did not call any of the “shuffled genomes” deemed to be false positives in the datasets. Compared to other programs presented in [18], StrainSeeker’s run time is one of the fastest and it also has the highest sensitivity. The results are available at http://bioinfo.ut.ee/strainseeker.

### Computational requirements and speed of the method

Using a UNIX server and 32 CPU cores, we created the database of 2,758 strains in approximately 8 hours, its size was 7.6 GB. Because StrainSeeker’s database building process creates some large temporary list files, the recommended disk space is at least 200 GB. If disk space is a constraint, the database can also be downloaded from our web page. To speed up the identification process, a large list of all k-mers present in the tree (the “whitelist”) is used to reduce the sample list size and keep only the k-mers that could be matched to the database.

A typical identification of a 140 Mbp *E. coli* isolate sample using the 2,758 strain database, UNIX server and a single CPU core took approximately 50 seconds. The identification of a large, 12.3 Gbp mock metagenome (SRA database, SRR2131179) took 26 minutes with StrainSeeker. The k-mer list creation step required the most RAM and was dependent on the FASTA/FASTQ file size. Because the assembled bacterial genomes are quite small, the minimum RAM required for database building is considerably less than 1 GB. However, sequenced samples with large amounts of reads require more RAM. For example, the identification of a 140 Mbp sample takes approximately 150 - 500 MB of RAM. StrainSeeker does not load the whole database into memory, which means that StrainSeeker could use very large databases and still work well even on personal computers. Similar programs [7, 8] require at least 4GB of RAM when using the smallest databases.

## Conclusions

There is a strong need for the fast detection of bacterial strains. StrainSeeker can detect previously unseen strains and provide a clear list of database strains closest to the unknown strain. Users can provide their own guide trees and build databases that contain their strains of interest. In the current study, we showed that StrainSeeker could obtain clear, strain-level identification results for samples that contained small amounts of an *E. coli* strain and large amounts of another *E. coli* strain. Minimum amount of reads required to detect the minor *E. coli* strain was 20,000 to 50,000. Also, StrainSeeker accurately identified strains in a diverse metagenomic sample even when human DNA was present (Additional data file 2). Still, due to the statistical framework which StrainSeeker uses, it has some limitations. First, it still requires several thousand reads to detect a strain, the exact amount varying with the genome size and other strains present in the sample. Second, it is not able to differentiate between strains that are distinguished by only a few single nucleotide variations. This requires high-quality assemblies which can be difficult and expensive to acquire for every sample. Third, only assembled genomes can be used as an input for StrainSeeker database building. However, the future updates/versions of the software should enable StrainSeeker to use raw, high-coverage reads from isolated strains. Initial tests indicated that adding high-coverage reads of a strain isolate to the database rather than adding an assembled genome gave better identification results. Moreover, assemblers often require fine-tuning of several parameters that can be cumbersome for non-bioinformaticians. All in all, StrainSeeker is a tool that can identify bacteria on the strain level from either isolates or mixed samples and provide clear, human readable output within minutes. There is also an online version of StrainSeeker freely available at http://bioinfo.ut.ee/strainseeker.

## Methods and materials

### *E. coli* isolation, DNA sequencing, genome assembly and initial identification

During a 5 month period in 2012. *E. coli* strains were isolated from samples taken from 21 hospitals located in Estonia (n=5), Latvia (n=4), Lithuania (n=3), Norway (n=1) and St. Petersburg (n=8). The strains were isolated from different clinical materials: blood, pus, urine and the respiratory tract. Initial bacterial identification was performed using MALDI-TOF MS (Maldi Biotyper, BrukerDaltonics GmbH, Germany). The ESBL phenotype was confirmed with the ESBL+AmpC identification kit (Rosco Diagnostica, Taastrup, Denmark). DNA templates for sequencing were generated by growing cultures of *E. coli* isolates overnight on blood agar (Oxoid Limited, UK). Total DNA from the bacterial strains was extracted using the QIAamp DNA Mini Kit (Qiagen, Germany). Bacterial genomic DNA was quantified using the Qubit® 2.0 Fluorometer (Invitrogen, Grand Island, NY, USA). A total of 1 ng of sample DNA was processed for the sequencing libraries using the Illumina Nextera XT sample preparation kit (Illumina, San Diego, CA, USA) according to the manufacturer’s instructions. The DNA normalization step was skipped; instead, the final dsDNA libraries were quantified with the Qubit® 2.0 Fluorometer and pooled in equimolar concentrations. The library pool was validated with 2200 TapeStation (Agilent Technologies, Santa Clara, CA, USA) measurements, and qPCR was performed with the Kapa Library Quantification Kit (Kapa Biosystems, Woburn, MA, USA) to optimize cluster generation. A total of 96 bacterial genomic libraries were sequenced with 2 x 101 bp paired-end reads on the HiSeq2500 rapid run flowcell (Illumina, San Diego, CA, USA). Demultiplexing was performed with CASAVA 1.8.2. (Illumina, San Diego, CA, USA) allowing 1 mismatch in the index reads. Genomes were assembled with the *de novo* assembly program Velvet [20].Prior to assembling, the reads were trimmed and filtered for quality (fastq_quality_trimmer–Q33–t 30–l 40, fastq_quality_filter–Q33–q 25–p 90) (http://hannolab.cshl.edu/fastx_toolkit/index.html). The cyclic assembly process was applied for each genome where different Velvet parameter values (*-exp_cov,-cov_cutoff (3, 5, 10, 15),-min_pair_count (1-5),-ins_length (100-350)*) were tested until all MLST genes were found or the best set of MLST genes was retrieved.

### Multi-locus sequence typing of E. coli samples

For accurate *E. coli* strain MLST type identification, we used the assembled *E. coli* genomes (described above) and a MLST tool published by Larsen and others [21] that calculates the MLST profile based on a BLAST alignment of the input sequence file and the specified allele set. Public *E. coli* database “#1” version 2014_01 for molecular typing was downloaded from PubMLST (http://www.pubmlst.org/).

### Building the guide tree

We used k-mer-based alignment-free methods analogous to [11, 12] to calculate the pairwise distance K between all pairs of genomes and to create the guide tree. All 2,758 available bacterial genomes from the NCBI RefSeq database (release 65) were used. For every two bacteria, the expected amount of shared k-mers (Eshared) was calculated and the observed amount (Oshared) was counted using the GenomeTester4 software [17]. The expected value was calculated by assuming their genome sequences were random strings. Additionally, we calculated the median M from all E_shared_/O_shared_ (E/O) ratios for all pairs. *E/O* < 1 indicates that the given bacteria have more common k-mers than expected from two random sequences of given lengths, suggesting they are most likely evolutionally related. We expect the numerical value to be proportional to the similarity of given strains, and thus it functions as the basis for pairwise distance in tree-building. To make E/O more linear for a wider range of phylogeny levels, we used the following formula to calculate the pairwise distance K: *K* = 2^*(E/O)/M*^. For each genome pair, three different K values were calculated with different k-mer lengths (16, 20 and 24); the arithmetic average of these lengths was used as the final value in the pairwise distance matrix. We constructed the guide tree with MEGA6 [16] using the derived distance matrix and UPGMA method.

### StrainSeeker identification algorithm and calculation of the relative genome frequencies

First, the algorithm converts sequencing reads to a k-mer list and maps the k-mers to the guide tree (Figure 4). The identification process starts at the root node and recursively moves down towards the leaves. For each step, the percentage of observed k-mers O is calculated for the current node N1: *O* = *N*1_*obs*_/*N1*_*tot*_*100%. *N*1_*obs*_ indicates N1-specific k-mers found in the sample and *N1*_*tot*_ indicates the total amount of k-mers at N1. If O is below a cutoff level (5%), the search will not continue below N1. Otherwise, an observed/expected ratio O/E is calculated for N1 with children C1 and C2 as follows: *O/E* = (*N*1_*obs*_/*N*1_*tot*_)/(*C*1_*obs*_/*C*1_*tot*_ + *C*2_*obs*_/*C*2_*tot*_ - *C*1_*obs*_/*C*1_*tot*_\**C*2_*obs*_/*C*2_*tot*_). *C*1_*obs*_ and *C*2_*obs*_ indicate the number of child-specific k-mers found in the sample and C1tot and *C2tot* indicate the total amount of child-specific k-mers. O/E shows whether less (*O/E* < 1, indicates mainly sequencing errors), more (*O/E* > 1, indicates that a new strain branches from the current node) or approximately the same amount (*O/E* = 1, indicates that at least one of the sub-nodes are present in the sample) of k-mers were found as expected if C1 and/or C2 were present in the sample. We used an asymptotic test with a significance level of 0.5*10^−3^ to test the hypothesis that *O/E* = 1 (additional data file 1). If we cannot reject the hypothesis, the search will continue until the strain level is reached with O and O/E calculated and checked at each step. If we reject the hypothesis, the search on the current branch will either stop (O/E < 1) or all strains under the current node will be reported in the output (O/E > 1). To calculate the relative genome frequencies, we assumed that the number of times a k-mer is seen follows Poisson distribution. To reduce the influence of possible errors (either due to sequencing errors or a k-mer not being unique), we ordered the k-mer list by frequency and removed both the top 10% and lowest 10% of the list. We calculated the mean coverage from the remaining k-mers. Based on the mean of truncated observations, the mean of non-truncated Poisson distribution is estimated using the maximum-likelihood estimation.

#### Creation of artificial test samples

18 *E. coli* samples were created by pooling together Illumina sequencing reads from each of the minor strains (*E. coli* ST73, ST131 and ST156 strain isolated from clinical samples as described above) and the major strain *E. coli* ST10 (*E. coli* K12 substrain MG1655; SRA database, run SRR892241). Each sample contained one million 100 bp reads. The major strain was used to add background noise in order to make the identification more difficult. The proportions of minor strains in the samples were 0.5%, 1%, 2%, 5%, 10% and 25% (Table 1).

To create the artificial metagenomic sample, we took a mock community metagenome sample from SRA database (SRR172902) and added raw human sequencing reads (SRA database, ERR1055492) to obtain a 1:10 ratio of bacterial DNA to human DNA. Human sequencing reads were trimmed to the same 75 bp length as the bacterial reads. Sample size was 4.92 Gbp.

## Software and data availability

StrainSeeker is written in PERL and is available for download or online use at [20] along with the sequencing reads of the three *E. coli* isolates used in the experiments.

## Additional data files

The following additional data are available with the online version of this paper. Additional data file 1 is a thorough description of the statistical test that is part of the StrainSeeker identification algorithm. Additional data file 2 contains StrainSeeker identification results of an artificial metagenomic sample (SRA database; SRR172902 mixed with ERR1055492).

## List of abbreviations

bp: base pair
NCBI: National Center for Biotechnology Information
MLST: multi-locus sequence typing
SRA: Sequence Read Archive
WGS: whole-genome sequencing
UPGMA: unweighted pair group method with arithmetic mean

## Authors’ contributions

MRo, MV, LK and ML wrote the software. MRo wrote the paper. PN and SK provided *E. coli* isolate data. MRe designed the experiments. TK performed the assembling of *E. coli* isolate sequencing reads. RA performed *E. coli* isolate MLST typing. MRo and MV performed the experiments and the analysis. MM designed the statistical framework. All authors read and approved the final manuscript.

## Acknowledgements

This work was supported by the European Union through the European Regional Development Fund (Estonian Centre of Excellence in Genomics and ARMMD project No 3.2.0701.11-0013), by the Estonian Ministry of Education and Research (institutional grant IUT34-11, target financing SF0180132s08 and KOGU-HUMB), by the Baltic Antibiotic Resistance collaborative Network (BARN), by the University of Tartu (SARMBARENG) and by the Estonian Science Foundation (grant No 9059).

